# Functional and phylogenomic approaches reveal novel types of M42 peptidases with contrasted enzymatic properties in Archaea

**DOI:** 10.1101/2025.02.26.640355

**Authors:** Emilie Chagny, Najwa Taib, Daphna Fenel, Eric Girard, Simonetta Gribaldo, Didier Flament, Bruno Franzetti

## Abstract

M42 peptidases are half-megadalton aminopeptidases characterized by a tetrahedral architecture (TET) ubiquitous across all domains of life. Despite their widespread occurrence, their evolutionary history and functional diversity remain largely unexplored. Here we show an unsuspected and largely untapped wealth of archaeal TET peptidases, exhibiting remarkable functional heterogeneity, as illustrated by the characterization of six novel enzymes. Using structural biology, phylogeny, and enzymatic studies, we establish robust criteria for high-throughput identification of TET peptidases and perform the first systematic study of their genomic distribution and functional diversity across the archaeal kingdom. We propose an 11-group classification for these enzymes and identify one group as the ancestral lineage that first emerged in Archaea. By coupling taxonomic distribution patterns with functional insights, we highlight the presence of multiple TET enzymes with selective activities in heterotrophic and mixotrophic organisms, suggesting a role for TET peptidases in the degradation of environmental peptides. Overall, this work illuminates the underexplored diversity of TET enzymes, uncovering a complex evolutionary history linked to their potential biological function.

## Introduction

The development of environmental surveys and culture-independent approaches, such as metagenomics and high-throughput sequencing, have greatly enhanced our understanding of the global microbial inventory, particularly in marine ecosystems^1–3^. This surge in genomic data availability unveiled an unforeseen level of taxonomic and functional diversity among prokaryotes, leading to the discovery of novel metabolic pathways, insights into enzyme functions related to environmental adaptation, and the identification of new biocatalysts for biotechnological applications^4,5^. However, functional annotation based on sequence or structural similarity, even with deep learning-based bioinformatics tools, often falls short of accurately classifying proteins^6^ and requires direct experimental validation. While high-throughput functional screening approaches have been used to identify enzymes of interest from genomic data^7^, these methods only allow exploration of a limited range of substrates and activation conditions. This is especially problematic for large complexes or extremophilic enzymes, which often require specific activation conditions. To address these challenges, we introduce an innovative hybrid approach integrating phylogenetic analysis with biochemical characterization, enabling the exploration of the functional diversity of M42 aminopeptidases, a unique type of giant self-compartmentalized aminopeptidases forming distinctive tetrahedral structures, named TET.

TET peptidases are known to sequentially degrade the N-terminal residues of peptides up to 40 amino-acids in length^8^. These enzymes belong to the M18 and M42 families of the MEROPS classification^9^. M18 members are predominantly found in eukaryotes and bacteria, while M42 peptidases are restricted to prokaryotes^10^. Although their biological significance remains poorly understood, they have been hypothesized to be involved in intracellular proteolysis downstream of the proteasome^8,11^, with no clear supporting evidence to date. Archaeal TET peptidases, in particular, have been studied extensively. Following the discovery of the first TET complex in *Haloarcula marismortui*^8^, several high-resolution structures of archaeal enzymes revealed a conserved architecture, characterized by the assembly of twelve subunits into a ∼450 kDa hollow tetrahedral complex. Dimers, which are the building block of the dodecamer, are positioned along the edges of the complex. The faces of the tetrahedron are defined by three dimers forming a central opening, which is presumed to function as the substrate entry site, leading to a wide inner cavity. All catalytic sites, situated in the apexes of the complex, are oriented toward this chamber^11–14^. The active site comprises seven catalytic residues, with five of these residues coordinating two metallic ion cofactors^12,15–17^.

Despite a high degree of structural conservation, characterization of several archaeal TETs revealed significant functional disparities, along with variations in the copy number per organism. The single TET of *H. marismortui* was characterized as a broad-spectrum aminopeptidase, whereas four homologous TETs exhibiting distinct and narrower substrate specificities were identified in *Pyrococcus horikoshii*: PhTET1, PhTET2, PhTET3, and PhTET4 are glutamyl-, leucyl-, lysyl-, and glycyl-specific aminopeptidases, respectively^13,18–21^. Owing to this functional versatility, TET peptidases hold significant potential for biotechnologial applications in nutrition, health, and cosmetics. For instance, archaeal TET peptidases can increase the diversity of bioactive peptides in hydrolysates derived from natural biomass, which could be used in the agri-food sector to improve the nutritional quality of feeds for aquaculture^22,23^. However, with only a few archaeal enzymes thoroughly functionally characterized so far^8,13,15,18–21,24^, mostly from closely related species within the Thermococcales order, it is uncertain whether current knowledge fully accounts for the functional diversity and biological activities of these enzymatic complexes.

In this study, we carried out a high-throughput screening for M42 TET peptidases in 3,702 archaeal genomes using structure-based identification criteria. We uncover a previously unknown diversity of TET spanning the whole tree of Archaea. By combining phylogenetic and biochemical analyses, we propose to classify archaeal TETs into eleven groups. Six new TETs from previously undescribed groups were characterized, revealing a large functional diversity of these enzymes. Finally, we infer the evolutionary history of TET peptidases and discuss new insights into their potential biological roles.

## Results

### Structure based analysis identifies a large diversity of TET peptidases in Archaea

Despite strong structural homology, M42 peptidase primary sequences exhibit high divergence^14^. This variation, coupled with frequent misannotations as cellulases or endoglucanases^24,25^, makes it challenging to identify M42 peptidases in genomic or proteomic databases by sequence homology searches. Using the structural elements involved in the formation of their unique tetrahedral architecture as key determinants, we identified several residues to better delineate M42 aminopeptidases (**Fig. 1**). First, the presence of the seven conserved catalytic residues (His62, Asp64, Asp173, Glu205, Glu206, Asp/Glu228, and His307 in PhTET1, PDB code 2WYR) coordinating two metallic cofactors is essential for both the catalytic activity and structural integrity of the complex^15–17^. These residues being shared by all peptidases of the MEROPS MH clan (*i.e.,* M18, M20, M28, and M42 families), additional criteria are needed for accurate M42 peptidase segregation^26^. Sequence and structure comparison of MH clan enzymes highlighted the unique presence of five glycine residues (Gly44, 77, 85, 86, 211 in PhTET1) in M42 peptidases that confer the required flexibility for proper protein folding in the periphery of the active site. Taken together, these two criteria provide a robust method for screening M42 aminopeptidases in genomic and proteomic databases. To ensure clarity and consistency with previous studies, the term ’TET peptidases’ will be used exclusively for the remainder of this article.

**Fig. 1:**
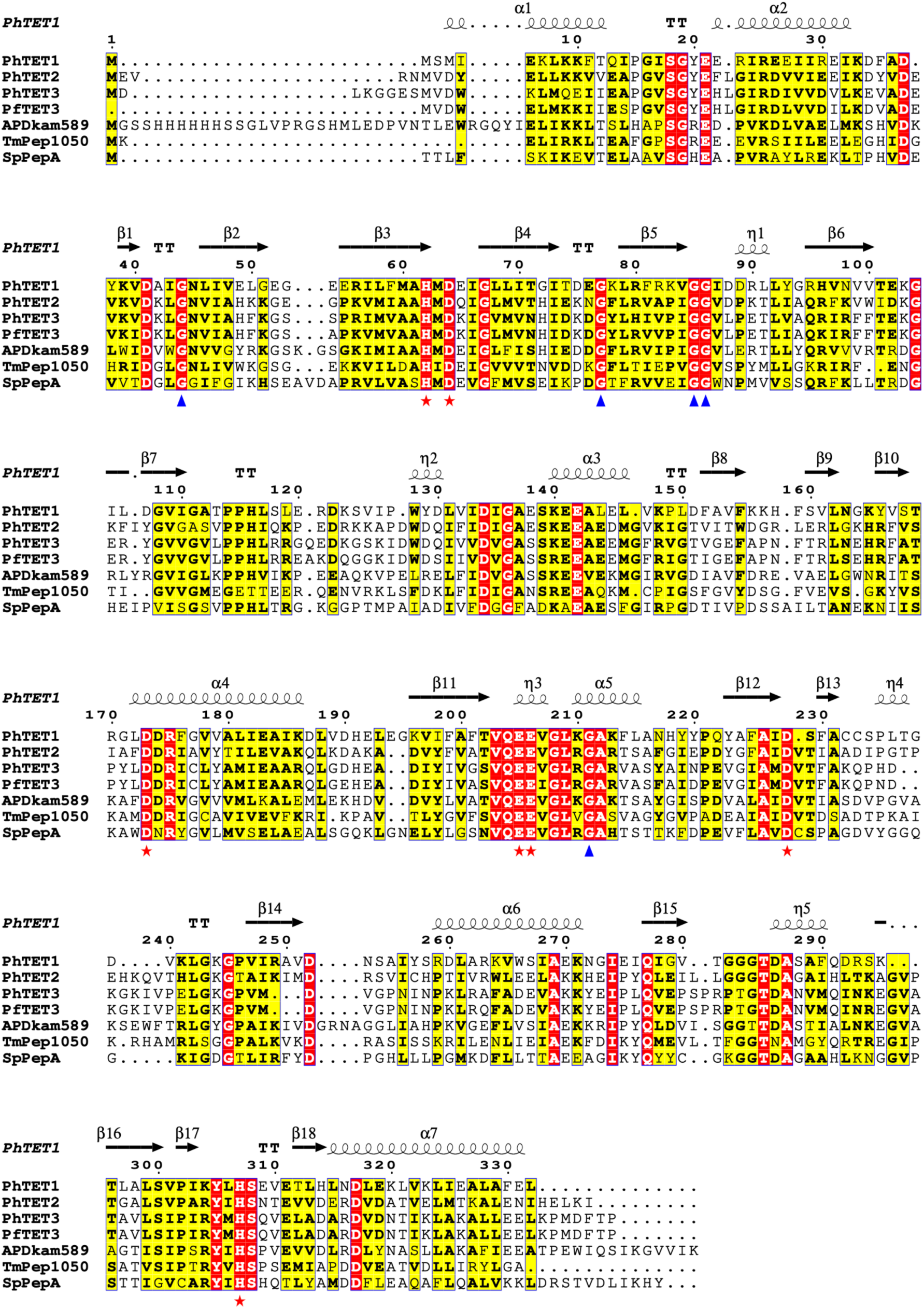
Structure-based determinants for identifying M42 peptidases. Multiple sequence alignment of structurally and functionally characterized M42 aminopeptidases. The represented secondary structure corresponds to PhTET1 (PDB code: 2WYR). Two criteria were retained for the identification of TET aminopeptidases: conserved catalytic residues H62, D64, D173, E205, E206, [DE]228, H307 (red stars), and conserved glycine residues 44, 77, 85, 86, and 211 (blue triangles). Residue numbering according to PhTET1 sequence.

An HMM profile was built using 210 archaeal TET sequences gathered from the MEROPS and NCBI nr databases. This profile was used to conduct an exhaustive homology search of TET peptidases against a large database containing 3,702 genomes of Archaea and covering all currently available diversity. All positive hits were retrieved and filtered based on the presence of the conserved catalytic site and glycine residues mentioned above, resulting in 1,826 archaeal TET homologues (**Supplementary Table 1**). TET peptidases were found in 39.0% of archaeal genomes (1,443) with varying numbers of TET copies per species, ranging from one to four. While TET homologues are widely present in Euryarchaeota and Asgard, their distribution is uneven to sporadic in the TACK and the DPANN superphyla, respectively (**Supplementary Table 1**). Intriguingly, some TET peptidase sequences from Asgard display a specific pattern consisting in an insertion of approximately 20 residues, which has never been detected in any TET peptidase described to date. This peculiar insertion, located just after the dimerization interface^21^, was found in 74 Asgard species and exhibits a strong charge contrast. A similar but shorter insertion (11-12 residues) was also observed in 23 species of Thermoplasmata (**Supplementary** Fig. 1).

We investigated the presence of possible functional analogs in genomes lacking TET peptidase. As previous studies suggested complementarity between M42 and M18 or TRI peptidases^11,25,27^, we used the PFAM domains PF02127 and PF14684 to search for M18 and TRI homologues in our local database of archaeal genomes (**Supplementary Table 1**). Our results challenge these hypotheses; M18 and TRI homologues were only sparsely detected, primarily in species possessing M42 peptidases. Furthermore, several lineages (*e.g.* Theionarchaea, Pontarchaeia, Thalassoarchaeia, Methanocellia, Thaumarchaeota) lack all three peptidase families. This points to a more complex relationship and suggests the existence of other functional analogs yet to be identified.

To understand how archaeal TET peptidases are related to each other, we inferred a maximum likelihood tree using the 1,826 archaeal M42 sequences (**Fig. 2**). Members of the TET1, TET2, TET3, and TET4 groups were already identified in Thermococcales species prior to this study, with several characterized representatives from *Pyrococcus horikoshii*, *Pyrococcus furiosus* and *Thermococcus onnurineus*^13,15,18–21,28,29^. In the TET phylogeny, these groups form four distinct and supported monophyletic clades (with UFB values of 100%). While TET2, TET3, and TET4 contain exclusively sequences from Thermococcales, TET1 also includes three sequences from Geoglobus, likely arising from a horizontal gene transfer (HGT) from Thermococcales to Archaeoglobales (**Fig. 2**).

**Fig. 2:**
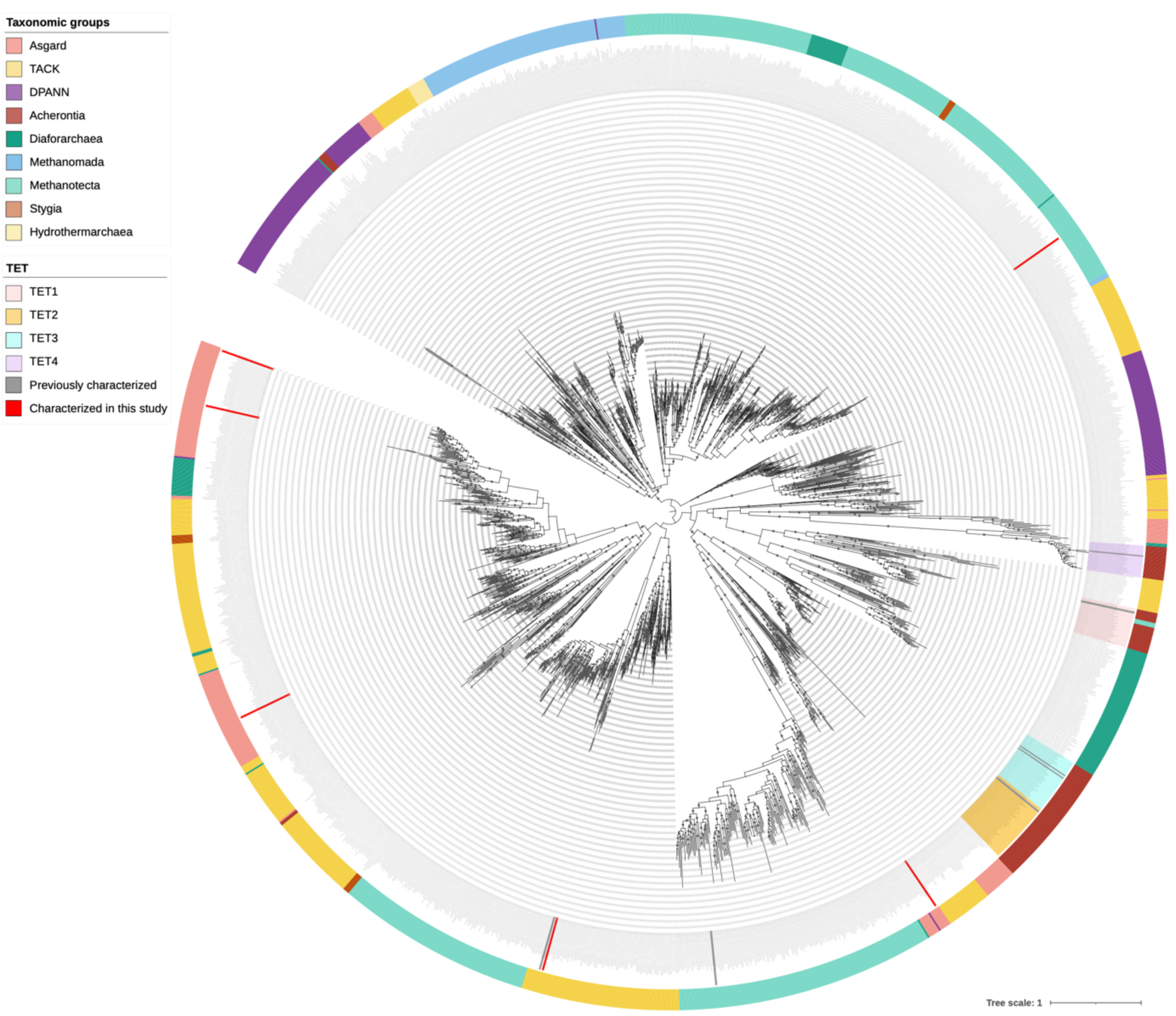
Phylogeny of archaeal TET peptidase homologues. Maximum-likelihood phylogeny obtained from an alignment of 1,826 sequences and 337 amino acid positions. The scale bar represents the average number of substitutions per site. Circles at the branches indicate ultra-fast bootstrap values >= 90 %. TET1 to TET4 families were delineated based on the taxonomic distribution and the topology of the tree. Gray and red bars on the inner circle indicate enzymes characterized prior to or during this study, respectively. Archaeal taxonomic groups are represented on the outer circle.

### Archaeal TET peptidases exhibit contrasting substrate specificities

Prior studies on the four TET of *P. horikoshii* reavealed distinct substrate specificities, already unveiling the functional versatility of these enzymes: PhTET2 functions as a broad-spectrum leucyl-aminopeptidase^18^, while PhTET1, PhTET3, and PhTET4 specifically target acidic, basic, and glycine residues, respectively^13,19,20^. However, a striking observation from the present analysis is that previous characterization of these few archaeal TET peptidases remains marginal with regard to the real taxonomic distribution and overall diversity of TET peptidases in archaeal genomes^8,13,15,18–20^. To fully explore their functional diversity in Archaea, we selected thirteen sequences, phylogenetically distant and from nine different species (1 TACK, 1 DPANN, 3 Asgardarchaeota, 2 Methanotecta, and 2 Methanomada) for recombinant protein expression and purification (**Table 1**, sequences provided in **Supplementary Table 2**). Notably, three Asgard sequences featuring the newly identified insertion were selected.

**Table 1:**
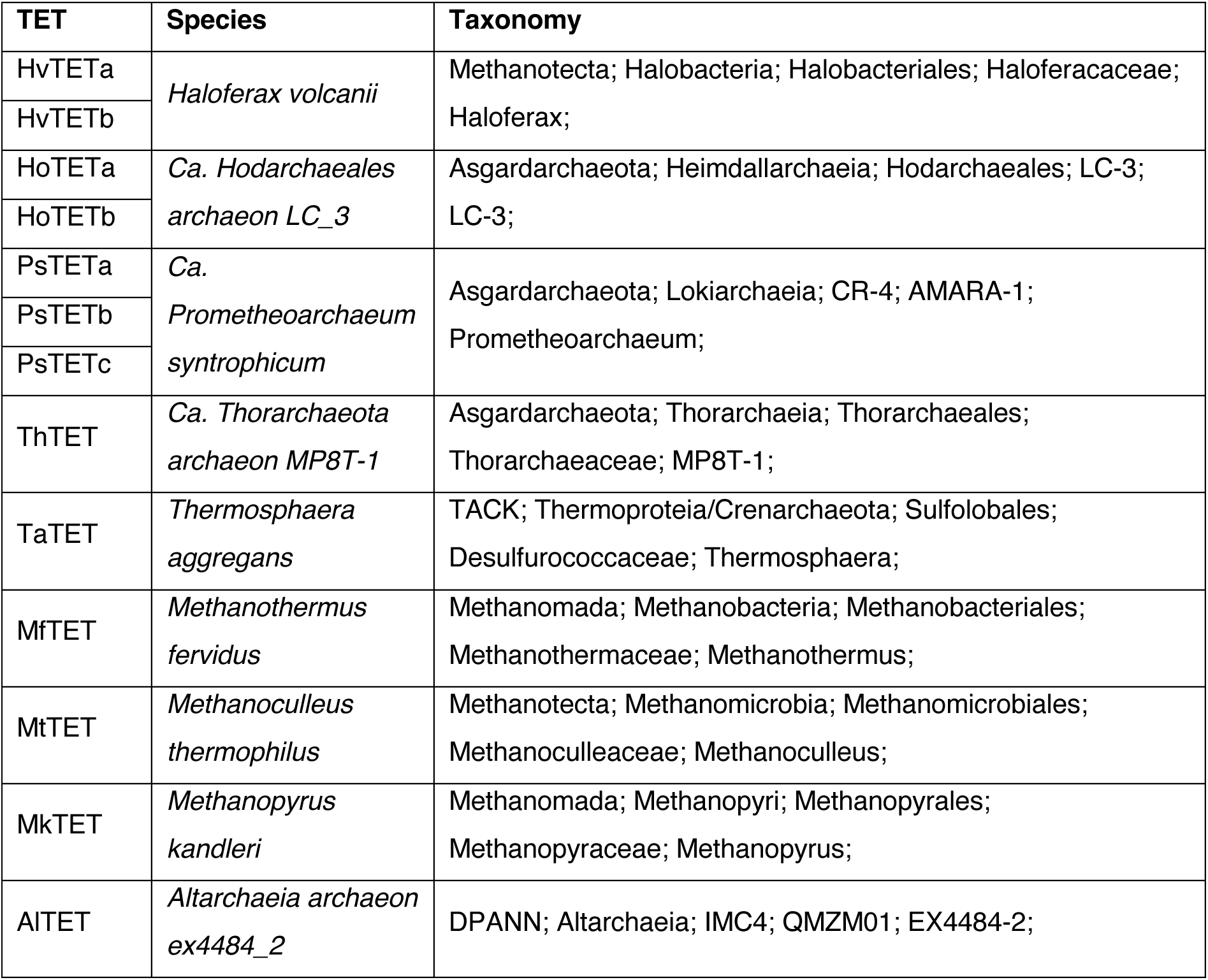
Candidate proteins for further characterization. Thirteen proteins spanning the phylogenetic tree and representing a broad range of taxonomically diverse species were selected.

Of the thirteen initially targeted proteins, six were successfully produced and purified to near homogeneity from *E. coli* extracts (*i.e.* HoTETb, PsTETa, PsTETc, ThTET, TaTET, and MtTET). Together with previously characterized TET peptidases, these new enzymes capture the core phylogenetic and taxonomic diversity of archaeal TETs. During the final gel filtration chromatography step, all proteins eluted as well-separated high molecular mass complexes corresponding to particles of a molecular mass of *c.* 450 kDa, indicating the formation of homo-dodecameric complexes (**Supplementary** Fig. 2). These findings were further validated by negative-stain electron microscopy observations, which revealed homogeneous populations of tetrahedral particles for most enzymes (**Supplementary** Fig. 2a). Lower purity of PsTETc and TaTET samples precluding satisfactory negative-staining imaging, structure predictions were generated using AlphaFold3 (**Supplementary** Fig. 2b). The resulting models displayed the expected hollow tetrahedral edifices with high confidence scores (ipTM 0.88 and 0.91), further supporting PsTETc and TaTET ability to form high molecular weight assemblies.

To investigate whether the diversity observed through phylogenetic analysis reflects functional diversity, cleavage specificities were studied using chromogenic [para-nitroaniline (pNA) conjugated] and fluorogenic [7-amino-4-methylcoumarin (AMC) conjugated] aminoacyl substrates (**Fig. 3**). The observed activity spectra are heterogenous and can be divided into two main clusters. PsTETa, ThTET, MtTET, and TaTET exhibit broad-spectrum activities and can be classified as generalist enzymes. These enzymes were found to preferentially cleave hydrophobic residues, with PsTETa and ThTET displaying broader specificities. PsTETa, ThTET, MtTET, and TaTET optimal amidolytic activities were observed with Ile-pNA, Leu-pNA, Met-pNA, and Met-pNA, respectively (**Fig. 3**). Similarities with the cleavage profile of PhTET2^18^ can be outlined, but PsTETa, MtTET, and TaTET represent the first description of methionyl and isoleucyl aminopeptidases in the TET family. Conversely, PsTETc and HoTETb exhibit more selective activities and can be classified as specialized enzymes. Analogous to the previously described PhTET1 peptidase, they specifically target acidic amino acids^19^. Interestingly, PsTETc maximum activity was measured on Glu-pNA, whereas no hydrolysis could be detected on Asp-pNA despite the similarity of these substrates (**Fig. 3**). The same substrate specificity has already been reported for the MHJ_0125 glutamyl-aminopeptidase of Mycoplasma hyopneumiae^30^. In addition to these varied substrate specificities, the characterized TET peptidases exhibit distinct activation profiles, with variations in optimal temperature, pH, and metal cofactor requirements (**Supplementary** Fig. 3).

**Fig. 3:**
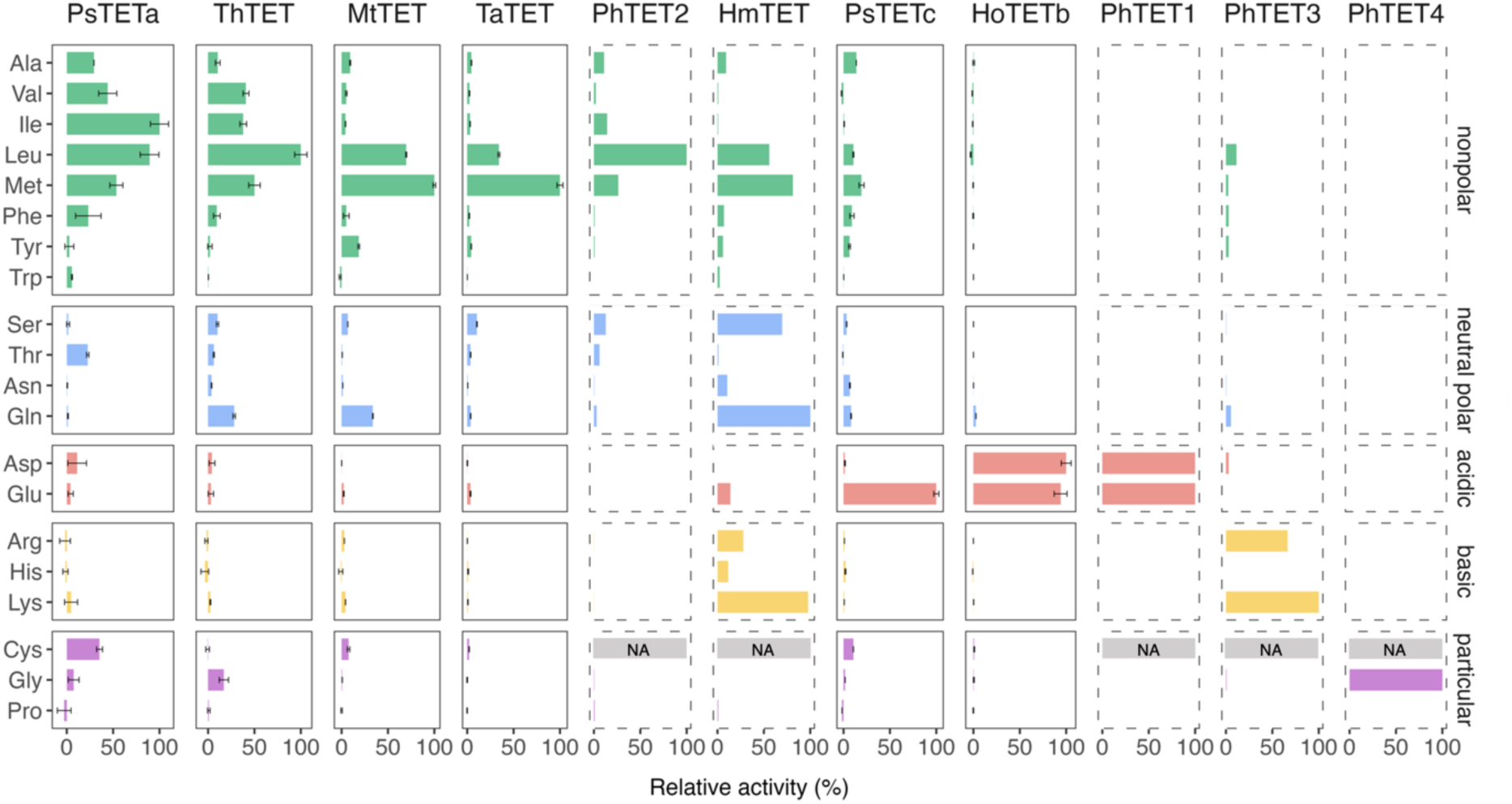
Characterized TET aminopeptidases exhibit diverse substrate specificities. Cleavage specificities were assayed using synthetic chromogenic and fluorogenic substrates. For each enzyme, activities are expressed as percentage of the maximum activity observed, which was attributed a value of 100%. Enzymes characterized prior to this study are indicated by dashed lines. Error bars indicate ±s.d. with n=3. NA: not assessed.

### Archaeal TET peptidases can be delineated into 11 groups

Considering that the pre-existing TET1 to TET4 groups cover only a small fraction of TET diversity, we used the distinct biochemical properties of the characterized enzymes and their taxonomic distribution to delineate new groups (**Fig. 3**). Specifically, contrasted substrate specificities led us to separate the TET6 and TET9 groups. Conversely, the broad TET11 group was maintained as a single group due to the similarities between the substrate specificities of the enzymes of *Methanocaldococcus jannaschii* (Atalah *et al.,* in preparation) and *Methanoculleus thermophilus*, along with a robust phylogenetic support. Similarly, TaTET has been identified as a generalist enzyme predominantly targeting hydrophobic residues, which is consistent with the substrate specificity of the previously characterized APDkam589 peptidase belonging to the same group^24^. Consequently, seven new groups were described across the tree, which were named TET5 to TET11 in accordance with the existing nomenclature (**Supplementary** Fig. 4), and each characterized enzyme was renamed according to its classification within the eleven defined groups. Notably, all sequences featuring the novel insertion described above were found in the TET7 group. Collectively, these eleven groups account for 1,500 sequences, the remaining 326 sequences were not affiliated to any group due to poorly supported branching or lack of characterized representatives.

Interestingly, TET11 emerges as the most prevalent group, distributed across the entire tree of Archaea, and is consistently present in methanogenic species, suggesting that this group may have been the first to appear in Archaea (**Fig. 4**). In contrast, TET1, TET2, TET3, and TET4 groups— previously the only known TET groups—appear to be restricted to Thermocci species (to the exception of a few TET1 sequences found in Archaeoglobales). TET2 and TET3 are sister clades, indicating that they arose from a gene duplication event. Interestingly, prior studies showed the *in vitro* and *in vivo* formation of PhTET2-PhTET3 heterocomplexes in *P. horikoshii*^16,31^. TET5 and TET6 members are exclusively detected in Halobacteria. TET7 and TET8 groups span two superphyla and are found in species from the Asgard and TACK groups. These sister clades also possess unique TET groups, with TET9 and TET10 being found exclusively in Heimdallarchaeia and Crenarchaeota, respectively (**Fig. 4**).

**Fig. 4:**
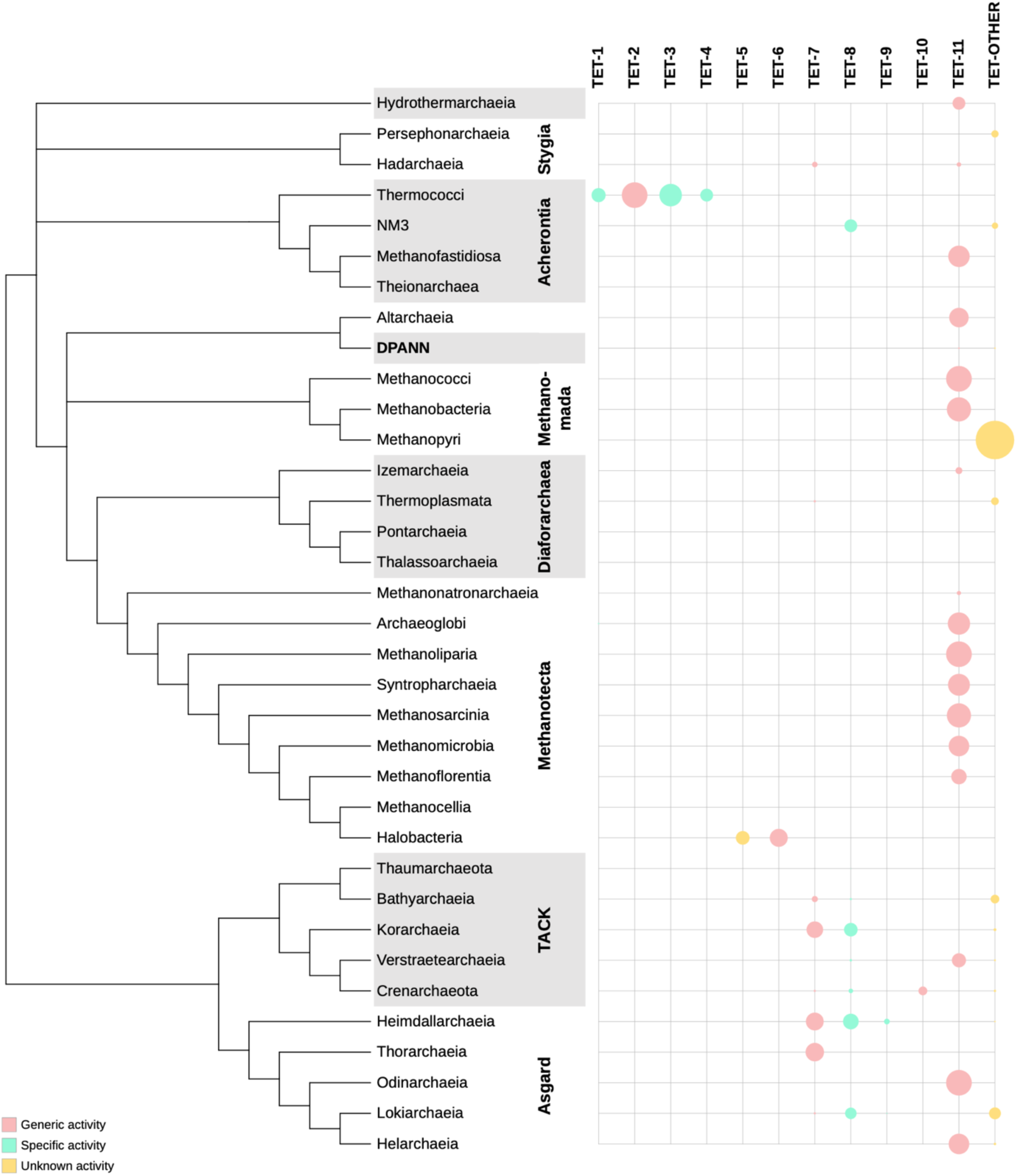
Phylogenetic distribution of the TET families in archaea. Distribution of the different TET families homologs on a schematic reference phylogeny of Archaea based on Garcia et al^49^. The sizes of the circles vary between 0% and 100% and indicate the percentage of genomes where a family is found. Circles are colored according to the activity spectrum of the characterized representatives of each family: pink for generic activities, green for specific activities, and yellow for undetermined activities

Finally, to investigate the origin of the TET peptidases, we searched for homologues in 401 representative bacterial genomes. We retrieved 187 bacterial TET homologues from 30% of the analyzed genomes displaying a patchy distribution across diverse phyla such as Thermotogae, Proteobacteria, Firmicutes, Chloroflexi, Deinococcota, Atribacteria, Bipolaricaulota, and Verrumicrobia (**Supplementary Table 1**). We inferred a maximum likelihood tree (**Supplementary** Fig. 5), which revealed a complex evolutionary history, shaped by multiple HGT intra and inter domains and several duplications. Two distinct groups can be delineated: the first group primarily contains archaeal sequences, spanning the full diversity of Archaea, with representatives from the TET1, TET4, and TET11 groups. Interestingly, sequences belonging to Bacteria, mainly Elusimicrobia, Thermotogae, Firmicutes, and Proteobacteria branch within this group indicating several independent HGT between archaea and these bacteria. The second group consists of a mixture of archaeal, mainly from the TACK and Asgard superphyla, and bacterial sequences, encompassing the remaining TET groups.

## Discussion

In this study, we used structure-based identification criteria for high-throughput screening of M42 peptidases to demonstrate a vast prevalence and diversity of these enzymes in archaea. Our approach notably revealed a ∼20-residue insertion, located close to the dimerization domain essential for TET particle assembly, in some Asgard TET peptidases. This insertion does not hinder oligomerization, since both PsTET7 and ThTET7 still form typical hollow tetrahedral particles. AlphaFold3 predictions suggest that these insertions may form protruding two-stranded β-sheets, potentially obstructing the substrate entry pores of the tetrahedral particle^21^ (**Supplementary** Fig. 6). Given that PsTET7 and ThTET7 demonstrate a broader substrate specificity relative to PhTET2, MtTET11, and TaTET10, this insertion may play a role in substrate recognition. Alternatively, this insertion might facilitate interactions with partner proteins. To fully elucidate the role of this novel insertion, further functional and structural studies on representative members of this group should be conducted.

Phylogenetic analysis and functional characterization of archaeal TET peptidases allowed us to propose a classification into eleven groups, seven of which had never been investigated before. An important finding from this study is the discovery of the TET11 group. These enzymes are the most prevalent and have nevertheless been completely overlooked until now. Indeed, while most of the TET groups are restricted to few taxa (*e.g.*, TET1-4 in Thermococci, TET5-6 in Halobacteria), the TET11 group is widespread in Archaea and is found in all superphyla, suggesting that it was present in the last archaeal common ancestor. It is also notable that species with a single TET enzyme tend to possess a member of the TET11 group. According to the characterization of MtTET11, TET11 members would display broad-spectrum activities. The ancestral origin of TET11, coupled with its enzymatic activity, suggest that this group was the first to appear in Archaea, followed by several duplications and horizontal transfers giving rise to multiple groups with different activities. Genetic studies on members of the proposed ancestral group TET11, such as the single TET peptidase of the genetically tractable species *Methanocaldococcus jannaschii*^32^, should be prioritized to establish the basic physiological role of the TET enzymes.

The study of TET co-occurrence within archaeal genomes showed that narrow substrate specificity occurs only in species harboring multiple TETs. This is the case for *P. horikoshii*, which possesses four TET peptidases of TET1, TET2, TET3, and TET4 groups. The four enzymes exhibit complementary activity spectra, suggesting that they function in concert to better achieve complete peptide hydrolysis^13,18–20^. Synergic specific activities of multiple TET peptidases have also been reported for the bacteria *Geobacillus stearothermophilus*^33^ and *Symbiobacterium thermophilum*^34^. Similarly, we identified specialized enzymes belonging to the TET8 and TET9 groups in the genomes of *Ca. Hodarchaeales archaeon LC_*3 and *Ca. P. syntrophicum* containing two and three putative TET peptidase genes, respectively. Multiplicity is also observed in other types of archaea such as Halobacteriales, which typically harbor both a TET5 and a TET6. HmTET6, the only characterized enzyme of the TET6 group, displays broad-spectrum activity. Although no enzyme from the TET5 group has been characterized to date, it could be hypothesized that this group exhibits narrower activity profiles. Accordingly, since TET-other group members are found in species possessing either a single TET or both a TET-other and an additional specialized TET, it can be hypothesized that enzymes display broad-spectrum activities.

Phylogenetic analysis indicates that TET multiplicity arose independently multiple times. This phenomenon is not exclusively attributable to HGTs, as evidenced by the emergence of the TET1 and TET4 groups by duplication within archaea. Contrasted substrate specificity and multiplicity of TET enzymes within archaeal proteomes does not appear to stem from environmental adaptation, as no correlation was identified between TET distribution and specific biotopes. For example, although Archaeoglobales, Thermococcales, Methanococcales, and Desulfurococcales all share the same ecological niche as primary colonizers of deep-sea hydrothermal vents^35–37^, these organisms exhibit markedly different TET distribution patterns. Conversely, the number and degree of specificity of TET peptidases present in an organism may correlate with its metabolic capabilities. Indeed, multiple TETs are found in heterotrophic and mixotrophic Hadarchaea^38^, Thermococcales^39–41^, Halobacteriales^42^, Crenarchaeota^43^ (*Vulcanisaeta* genus), Heimdallarchaeota^44–46^, Korarchaeota^47,48^, and Bathyarchaeota^49–51^ species. This suggests that TETs may play a metabolic role in the degradation of environmental peptides used as carbon sources, enabling efficient organic matter utilization. In contrast, TET peptidases are typically found in single copy in autotrophic species such as methanogens, which do not depend on the degradation of exogenous peptides, hinting at an alternative physiological role for these enzymes. As initially proposed, TETs may participate in protein homeostasis and amino acid recycling by processing peptides downstream of the proteasome and other related proteolytic complexes^8,11^.

To conclude, the hybrid approach adopted in this study—integrating structural biology, phylogeny, and biochemistry—revealed an unsuspected diversity of TET peptidases. This strategy shed light on a complex evolutionary history, uncovering an ancient subgroup of archaeal enzymes that had so far gone unnoticed. Moreover, we could classify archaeal TET peptidases into eleven distinct groups. Characterization of representatives from these groups revealed contrasting biochemical properties, underscoring the value of this approach to facilitate the discovery of proteins with discriminating characteristics within the same enzymatic family.

## Material and methods

### M42 peptidases identification in bacterial and archaeal genomes

To study the taxonomic distribution and the evolution of the M42 peptidase family in Archaea, we assembled a large database containing 3,702 archaeal genomes and 401 bacterial genomes representatives of all major phyla available in public databases as of January 2022 (**Supplementary Table 1**).

For homology searches, we built a specific HMM profile for the M42 archaeal peptidase family. For this, we used the MEROPS database (v12.0)^9^ and retrieved all archaeal M42 peptidase sequences longer than 260 amino acids (195 sequences). We estimated TET4 homologues to be under-represented in this dataset so in parallel, the LD[AE][EL]EKKED pattern, canonical for the TET4 group, was used to search the National Center for Biotechnology Information (NCBI) nr database restricted to Archaea using PHI-BLAST (default parameters)^52^. We retrieved 43 hits and used the T-Coffee trim tool (v11.0.8)^53^ to identify the 15 more divergent sequences, which were added to the initial set. Finally, the 210 resulting sequences were aligned with T-Coffee using default parameters, and the alignment was used to build an HMM profile using the HMMBUILD tool from the HMMER suite (v3.3.2)^54^.

This profile was used to carry out homology-based searches against our local Archaea database using HMMSEARCH. All hits were retrieved, aligned using MAFFT (v7.481, with the option -auto)^55^ and filtered upon the presence of the conserved motifs characterizing M42 peptidases (*i.e.*, residues Gly44, His62, Asp64, Gly77, Gly85, Gly86, Asp173, Gl205, Glu206, Gly211, Asp/Glu228 and His307 according to PhTET1 numbering, PDB code 2WYR). This resulted in 1,826 archaeal M42 peptidase homologues (**Supplementary Table 1**). These sequences were aligned using MAFFT (with the option -linsi), and the resulting alignment was trimmed using trimAl (v1.5.0, with the option -gappy-out)^56^. Finally, a maximum likelihood phylogeny was inferred using IQ-TREE (v2.0.6)^57^ with the model LG+F+R10 selected by ModelFinder according to BIC criteria^58^.

To investigate the origin and evolution of archaeal M42 peptidase, we extracted a reduced and taxonomically balanced database covering both archaea and bacteria from our local database (593 Archaea and 401 Bacteria, see **Supplementary Table 1**). Homology searches, alignment, and filtering steps were conducted as described above, yielding in 339 archaeal and 187 bacterial M42 peptidase homologues (**Supplementary Table 1**). The 526 sequences were aligned using MAFFT (with the option -linsi), and trimmed using trimAl (v1.5.0, with the option -gappy-out)^56^. A maximum likelihood phylogenetic tree was inferred using IQ-TREE and the model LG+R10 selected by ModelFinder according to BIC criteria (**Supplementary** Fig. 5). All phylogenies were annotated using IToL^59^.

Finally, we used HMMSEARCH (with the option -cut_nc) and the PFAM domains PF02127 and PF14684 to search for peptidases M18, and TRI, respectively, in our local database of Archaea (**Supplementary Table 1**).

### Bacterial strains and general information

*Escherichia coli* DH5α and Rosetta 2(DE3)pLysS chemically competent cells were used for cloning and recombinant expression, respectively. Cells were grown in lysogeny broth (LB) media in a rotary shaker at 37°C (or 20°C when specified), 140 rpm. When used, final concentrations of kanamycin and chloramphenicol were 30 μg/mL and 34 μg/mL, respectively.

For SDS-PAGE analysis, protein samples were mixed with loading buffer (50 mM Tris-HCl, 8 M urea, 2 M thiourea, 75 mM DTT, 3% SDS, 0.05% bromophenol blue, pH 6.8) in a 1:3 ratio, heated to 100°C for 4 min, and loaded on 12% Criterion^TM^ XT Bis-Tris Protein gels (BioRad). Protein bands were visualized by staining with InstantBlue (Expedon). Molecular weights were estimated relative to Precision Plus Protein All Blue Prestained Standards (Biorad).

### Expression and purification

The open reading frames of the selected genes were optimized for *E. coli* codon usage and synthesized by Twist Bioscience. For tagged protein expression, synthetic genes were digested with *NdeI* and *BamHI* restriction enzymes and inserted into the pET28a(+) vector, in frame with a thrombin-cleavable N-terminal His6-tag. For untagged protein expression, genes were digested with *NdeI* and *XhoI* restriction enzymes and cloned into the pET41c(+) vector. Cloning accuracy was assessed by Sanger sequencing (Eurofins).

The resulting recombinant plasmids were used for transformation of *E. coli* Rosetta 2(DE3)pLysS cells according to standard procedures^60^. Overnight cultures were diluted 1:100 and grown at 37°C, 140 rpm until OD_600_ reached 0.6. Protein overexpression was induced with 1 mM of isopropyl-β-D-thiogalactopyranoside (IPTG) for 16 h at 20°C. Cells were harvested by centrifugation at 8,000 × *g* for 45 min at 4°C, and pellets were stored at -80°C. Cells were resuspended in lysis buffer (50 mM Tris-HCl, 150 mM NaCl, 0.1% Triton ×100, pH 8.0) supplemented with 0.05 mg/mL lysozyme, 0.01 mL/mL MgSO_4_ 2M, 1 mg/mL Pefabloc SC, 0.05 mg/mL DNase, 0.2 mg/mL RNase, and were disrupted on ice in a Vibra-Cell sonifier (35% amplitude with five on/off cycles of 30 s each). For thermostable proteins, the lysate was heated at 70°C for 15 min. Insoluble particles were pelleted by centrifugation (16,000 × *g* for 30 min at 4°C) and the cleared extract was filtered at 0.45 μm and 0.22 μm. The recombinant proteins were purified from the soluble fractions to near homogeneity using various combinations of affinity, anion exchange and gel filtration chromatography.

For ThTET purification, after cell lysis, incubation at 70 °C for 15 min and clarification, the resulting supernatant was supplemented with imidazole (final concentration 10 mM) and loaded on a HiTrap Chelating HP 5 mL column (Cityva) equilibrated with 50 mM Tris-HCl, 150 mM NaCl, 10 mM imidazole, pH 8.0. Bound proteins were eluted with a linear gradient of imidazole (10 to 500 mM). Fractions corresponding to the elution peak at 400 mM imidazole were pooled, dialysed against 50 mM Tris-HCl, 20 mM NaCl, pH 8.0 and loaded on a ResourceQ column (Cytiva) equilibrated with the same buffer. Elution was achieved by a linear NaCl gradient (20 to 500 mM) and fractions containing protein of similar mass (37-39 kDa) according to SDS-PAGE were combined and concentrated using an Amicon Ultra-15 ultrafiltration unit (Millipore) with a 30 kDa cutoff. The protein was utlimately loaded on a Superose 6 Increase 10/300 GL column (Cytiva) in 50 mM Tris, 150 mM NaCl, pH 8.0. Fractions from the elution peak corresponding to a molecular mass around 450 kDa were pooled and subsequently concentrated using an an Amicon Ultra-15 ultrafiltration unit (Millipore) with a 30 kDa cutoff.

For HoTETb purification, after cell lysis, incubation at 70 °C for 15 min and clarification, the resulting supernatant was diluted to a final NaCl concentration of 75 mM and loaded on a ResourceQ column (Cytiva) equilibrated with 50 mM Tris-HCl, 75 mM NaCl, pH 8.0. Elution was achieved by a linear NaCl gradient (75 to 300 mM) and fractions containing protein of similar mass (37-39 kDa) according to SDS-PAGE were combined and concentrated using an Amicon Ultra-15 ultrafiltration unit (Millipore) with a 30 kDa cutoff. The protein was then loaded on a Superose 6 Increase 10/300 GL column (Cytiva) in 50 mM Tris-HCl, 150 mM NaCl, pH 8.0. Fractions from the elution peak corresponding to a molecular mass around 450 kDa were pooled and subsequently concentrated using an Amicon Ultra-15 ultrafiltration unit (Millipore) with a 30 kDa cutoff.

For TaTET purification, after cell lysis, incubation at 70 °C for 15 min and clarification, the resulting supernatant was dialysed against 50 mM Tris-HCl, 50 mM NaCl, pH 8.0 and loaded on a ResourceQ column (Cytiva) equilibrated with the same buffer. Elution was achieved by a linear NaCl gradient (50 mM to 1 M) and fractions containing protein of similar mass (37-39 kDa) according to SDS-PAGE were combined and concentrated using an Amicon Ultra-15 ultrafiltration unit (Millipore) with a 30 kDa cutoff. The protein was then loaded on a Superdex 200 10/300 GL column (Cytiva) in 50 mM Tris-HCl, 150 mM NaCl, pH 8.0. Fractions from the elution peak corresponding to a molecular mass around 450 kDa were pooled and subsequently concentrated using an Amicon Ultra-15 ultrafiltration unit with a 30 kDa cutoff.

For PsTETa, PsTETc, and MtTET purification, after cell lysis and clarification, the resulting supernatant was dialysed against 50 mM Tris-HCl, 20 mM NaCl, pH 8.0 and loaded on a DEAE sepharose CL-6B resin (Cytiva, XK16/20 column) equilibrated with the same buffer. Elution was achieved by a linear NaCl gradient (20 to 600 mM) and fractions containing protein of similar mass (37-39 kDa) according to SDS-PAGE were combined, dialysed against 50 mM Tris-HCl, 50 mM NaCl, pH 8.0 and loaded on a ResourceQ column (Cytiva) equilibrated with the same buffer. Elution was achieved by a linear NaCl gradient (50 to 500 mM) and fractions containing the protein of interest were pooled and concentrated using an Amicon Ultra-15 ultrafiltration unit (Millipore) with a 30 kDa cutoff. The protein was then loaded on a Superdex 200 10/300 GL column (Cytiva) in 50 mM Tris-HCl, 150 mM NaCl, pH 8.0. Fractions from the elution peak corresponding to a molecular mass around 450 kDa were combined and subsequently concentrated using an Amicon Ultra-15 ultrafiltration unit with a 30 kDa cutoff.

### Negative-stain electron microscopy

4 µL of purified protein samples (0.1 mg/mL) were absorbed onto the clean side of a carbon film on mica, stained, and transferred to a 400-mesh copper grid. Images were taken under low dose conditions (<10 e^-^/Å^2^) with defocus values between 1.2 and 2.5 μm on a Tecnai 12 LaB6 electron microscope at 120 kV accelerating voltage using CCD Camera Gatan Orius 1000.

### AlphaFold model predictions

AlphaFold model predictions were calculated using the AlphaFold3 server (https://alphafoldserver.com/ accessed on May 17^th^, 2024).

### Enzymatic characterization protocol

M42 peptidase hydrolytic activities on synthetic chromogenic and fluorogenic substrates were assayed using aminoacyl-para-nitroaniline (pNA) and aminoacyl-7-amino-4-methylcoumarin (AMC) conjugates ordered from Bachem. Substrates were solubilized in 100% dimethylsulfoxide (DMSO) to a final concentration of 20 mM. All assays described below were carried out according to the following standard procedure^18^. Reactions were initiated by addition of 2 to 10 µg/mL of enzyme to a pre-warmed mixture containing 2.5 mM of the synthetic substrate in 50 mM buffer (pH 5,5 – 11), 150 mM KCl, and 1 mM XCl_2_ (X = Ca, Co, Fe, Mg, Mn, Ni or Zn) in a total volume of 60 μL. To avoid water evaporation, the total volume was covered by 25 μL of mineral oil. Incubations were performed for 3 min to 1 h, reactions were stopped by the addition of 60 μL of 0.1 M acetic acid, and samples were placed on ice. After centrifugation at 6,000 × *g* for 3 min, liberated pNA or AMC quantities were quantified by OD_405_ or fluorescence (excitation and emission wavelengths 360 nm and 460 nm, respectively) measurement in a Synergy HT microplate reader (BioTek). Three replicates and two enzyme blanks were assayed for each experimental point. Enzyme concentrations and incubation durations were adjusted for each peptidase to produce a robust signal for accurate measurement.

For each enzyme, optimal temperature, pH and metallic cofactor were determined using the substrate on which maximum activity was measured. The effect of temperature on M42 peptidase activities was evaluated between 20 and 100°C. Assays were conducted as previously described in presence of 50 mM HEPES, 150 mM KCl, and 1 mM CoCl_2_, pH 7.5. To prevent enzyme denaturation and to ensure stable enzymatic activity, optimal pH and metallic cofactor were established 10°C below the determined optimal temperature. The effect of metal cations on M42 peptidase activities was assessed using 1 mM (0.1 mM for TaTET) of XCl_2_ metal (X = Ca, Co, Fe, Mg, Mn, Ni or Zn) with 50 mM HEPES, 150 mM KCl, pH 7.5. The influence of pH was studied in presence of 1 mM CoCl_2_ (0.1 mM for TaTET) using the following buffers: MES (pH 5.5 to 6.5), HEPES (pH 7.0 to 8.0), CHES (pH 8.5 to 9.5), and CAPS (pH 10.0 to 11.0). For each peptidase, substrate specificity was determined using optimal metal cofactor and pH. Incubation was performed 10°C the established optimal temperature.

| **TET** | **Enzyme concentration** | **Incubation duration** |
| --- | --- | --- |
| **HoTETb** | 2 μg/mL | 3 min |
| **MtTET** | 8 μg/mL | 10 min |
| **PsTETa** | 10 μg/mL | 60 min |
| **PsTETc** | 10 μg/mL | 20 min |
| **TaTET** | 10 μg/mL | 15 min |
| **ThTET** | 10 μg/mL | 3 min |

## Data availability

Data used to produce our results are provided as supporting data and can be found here: https://data.mendeley.com/preview/rc9nntwy6b?a=d335eeda-c9b3-429e-ae60-c07f432bf7cf.

## Acknowledgments

We thank Audrey Bossé for her technical support throughout this project. This work used the platforms of the Grenoble Instruct-ERIC center (ISBG ; UAR 3518 CNRS-CEA-UGA-EMBL) within the Grenoble Partnership for Structural Biology (PSB), supported by FRISBI (ANR-10-INBS-0005-02) and GRAL, financed within the University Grenoble Alpes graduate school (Ecoles Universitaires de Recherche) CBH-EUR-GS (ANR-17-EURE-0003). IBS acknowledges integration into the Interdisciplinary Research Institute of Grenoble (IRIG, CEA). This work was financially supported by the Région Bretagne, the French Research Institute for Exploitation of the Sea, and the Grenoble Alliance for Integrated Structural Cell Biology (GRAL). This work was supported by the Bettencourt-Schueller Foundation programme Impulscience (ENVOL) to S.G., Laboratoire d’Excellence ‘Integrative Biology of Emerging Infectious Diseases’ (grant no. ANR-10-LABX-62-IBEID), and the Fondation pour la Recherche Médicale (FRM). This work used the computational and storage services (TARS cluster) provided by the IT department at Institut Pasteur, Paris.

## Notes

https://data.mendeley.com/preview/rc9nntwy6b?a=d335eeda-c9b3-429e-ae60-c07f432bf7cf

### Competing Interest Statement

The authors have declared no competing interest.

