## Supplementary Figures for "Functional and phylogenomic approaches reveal novel types of M42 peptidases with contrasted enzymatic properties in Archaea"

***Supplementary Fig. 1: Multiple sequence alignment of 11 archaeal M42 peptidases illustrating a previously unreported insertion in Asgard and Thermoplasmata sequences.*** *The unique ~20 residue insertion, specific to some Asgard species, is highlighted in orange. A shorter but similar insertion (9-10 residues), highlighted in green, was identified in certain Thermoplasmata sequences.*

```
                                               150       160       170                             180
Phorikoshii@BAA30637.1                   ESKEEAEDMGVKIGTVITWDGRLERL.............GK.......HRFVSIAFDDRI
Phorikoshii@BAA30940.1                   SSKEEAEEMGFRVGTVGEFAPNFTRL.............NE.......HRFATPYLDDRI
Gimiplasmataarc@GCA020725925.1_00987     SNEEEAKAMGVRIGNPVVPDSCFSTMQKRIF.......KDGK.KSGSDTIAIGKAFDDRI
Gimiplasmatalesarc@UCE91752.1            SNEEEAKAMGVRIGNPVMPDSLFSTMTKRVF.......RDGK.RKGSDTIAIGKAFDDRI
Heimdallarcharc@GCA018238585.1_00239     SNAEEAKELGIRIGDPIAPHSIFEIWERPRIVKG.KD..GKEDEKSTVKLAVGKAFDDRA
Hodarchaealearc@GCA015520535.1_01757     KNKEEVEALGIQIGDPIVPDSKFELITRKEIIKD.SD...GKKQEKDVTLAMGKAFDDRI
Borrarchaealesarc@NHK31695.1             KSKKEVEALGIKLGDPAVPDSTFELLKRTQIIKD.ED..TKKKTKKEVTLAIAKGFDDRI
Thorarchaeacarc@GCA002825515.1_01215     SSAKEVKDLGIRVGDPAVPVASFRTLKRTRKEKKNEEDMDSKEETREVTLAVAKAFDDRI
Lokiarchaeiaarc@MBN2157246.1             KSDKEVKDLGIRIGDVASQYAIYRTMDRTRFKKD.DD...GKETKSTTKIAVAKAFDDRI
Phorikoshii@BAA29607.1                   ESKEEALEL.VKPLDFAVFKKHFSVL.............NG.......KYVSTRGLDDRF
Phorikoshii@BAA29828.1                   EKREDTEKLGIRPGDFIAFDPKFEYV.............N........GFVKSHFLDDKA
```

***Supplementary Fig. 2: New TET peptidases purification.*** *Size exclusion chromatography purification of ThTET and HoTETb was carried out using a Superose 6 Increase 10/300 GL column. Other proteins were purified on a Superdex 200 10/300 GL column. All proteins eluted as high molecular mass complexes of c. 450 kDa. (a) Negative-stain electron microscopy observation of ThTET, HoTET, PsTETa, and MtTET revealed homogenous populations of hollow tetrahedral particles. (b) Typical TET structures were predicted for PsTETc (blue) and TaTET (pink) using AlphaFold3 (ipTM 0.88 and 0.91, respectively). Models were aligned with PhTET2 structure (PDB code 1Y0R, in green), with RMSD of 0.783 Å for PsTETc and 0.534 Å for TaTET.*

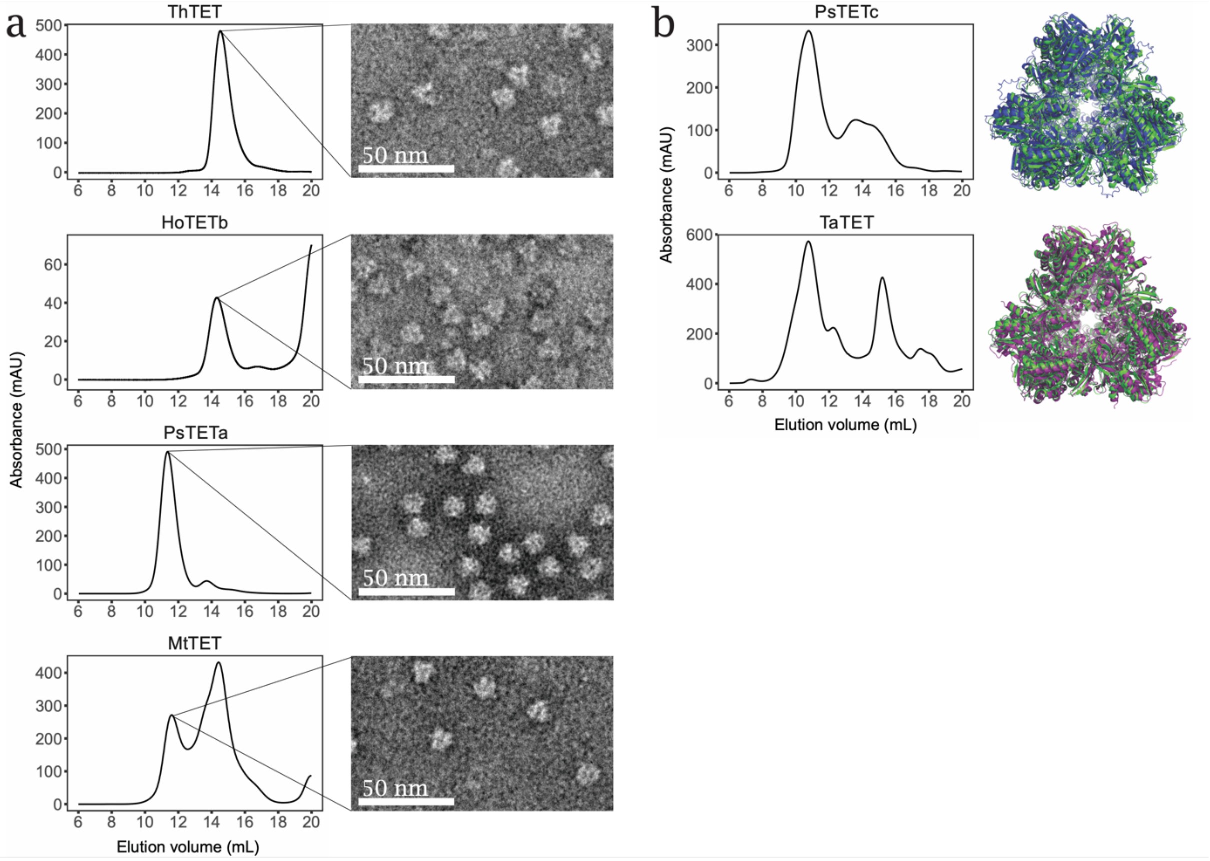

***Supplementary Fig. 3:*** *Temperature, pH and metal ions influence on TET enzymatic activities. For each enzyme, optimal conditions regarding temperature (top), pH (middle) and divalent cations (bottom) were determined. Error bars indicate ±s.d. with n=3.*

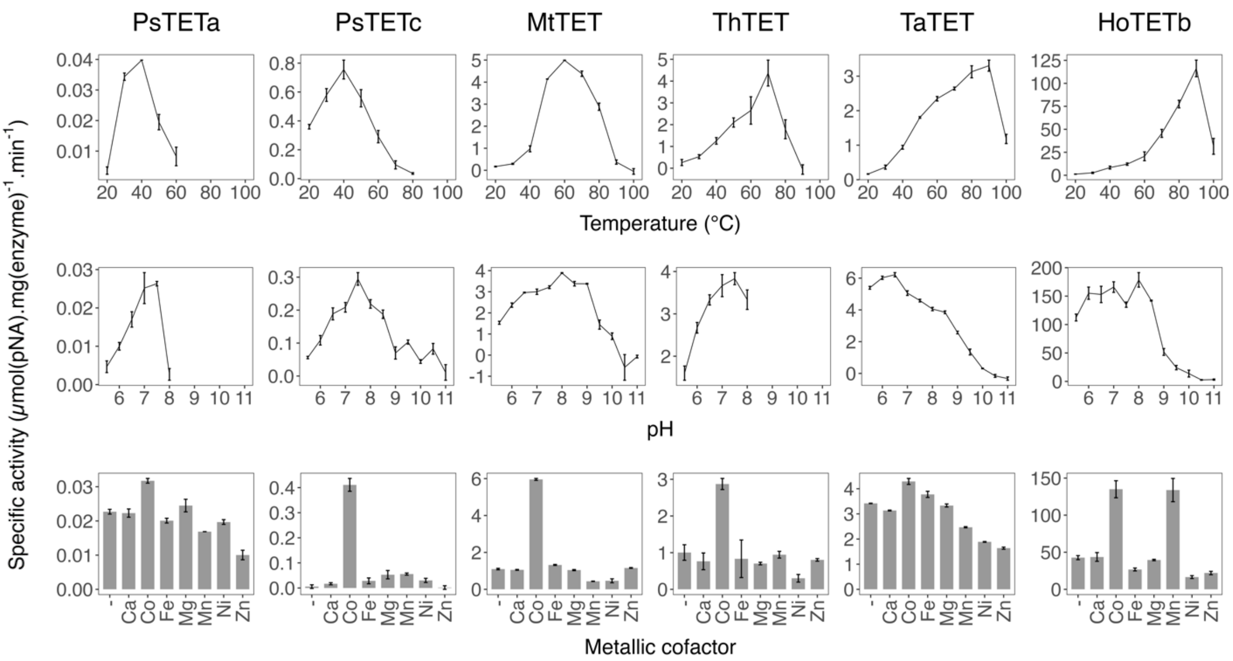

***Supplementary Fig. 4: Delineation of TET peptidases families in Archaea.*** *Maximum-likelihood phylogeny obtained from an alignment of 1,826 sequences and 337 amino acid positions. The scale bar represents the average number of substitutions per site. Circles at the branches indicate ultra-fast bootstrap values >= 90 %. TET families were delineated based on the taxonomic distribution, the topology of the tree (i.e. branch lengths, node supports) and substrate specificities. Gray and red bars indicate enzymes characterized prior to or during this study, respectively.*

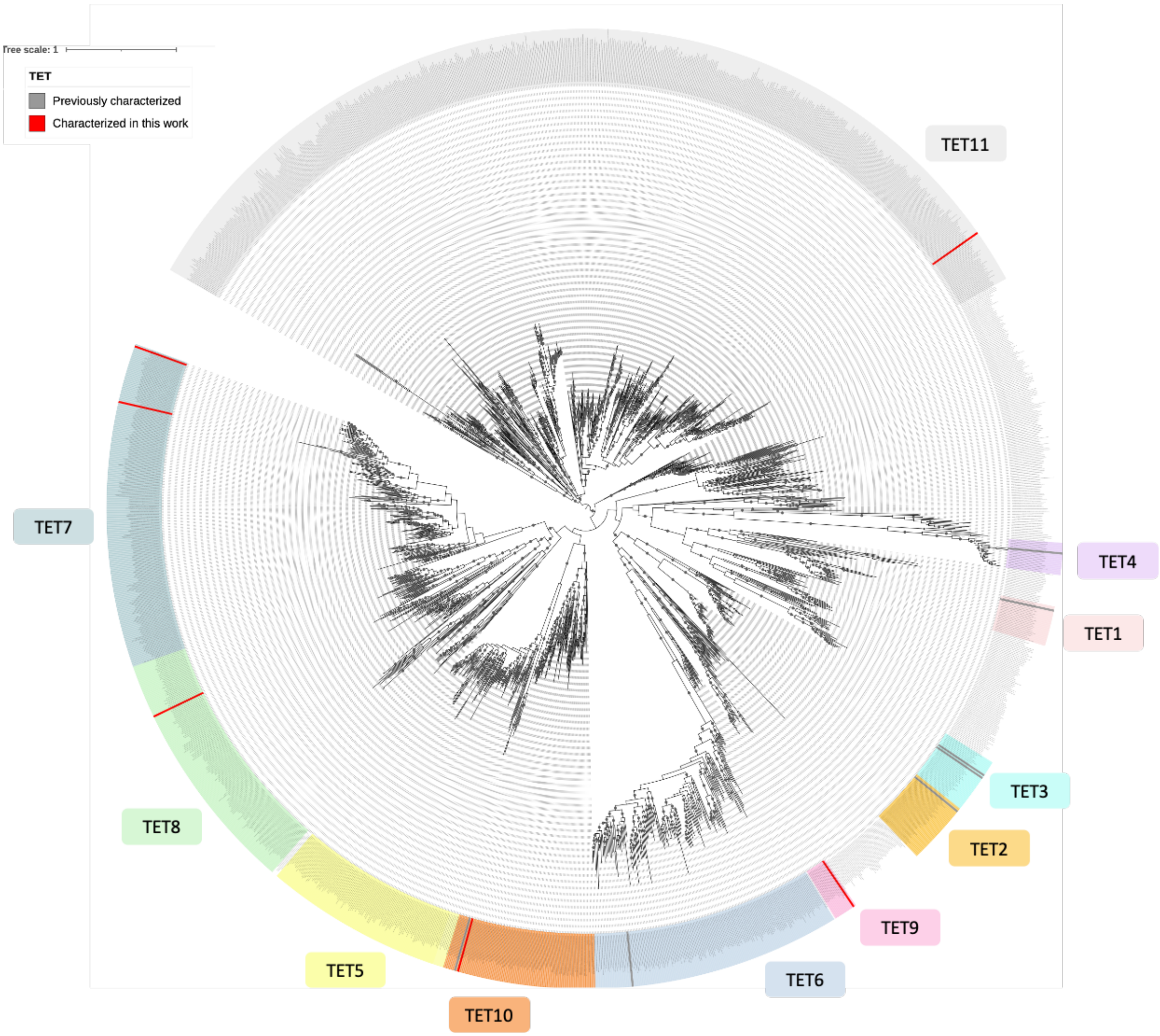

***Supplementary Fig. 5: Phylogeny of archaeal and bacterial M42 peptidase homologues.*** *Maximum-likelihood phylogeny obtained from an alignment of 526 sequences and 337 amino acid positions. The scale bar represents the average number of substitutions per site. Circles at the branches indicate ultra-fast bootstrap values ≥ 90 %. Archaeal and bacterial sequences are indicated in red and blue, respectively. The eleven group-classification for archaeal TET peptidases is represented.*

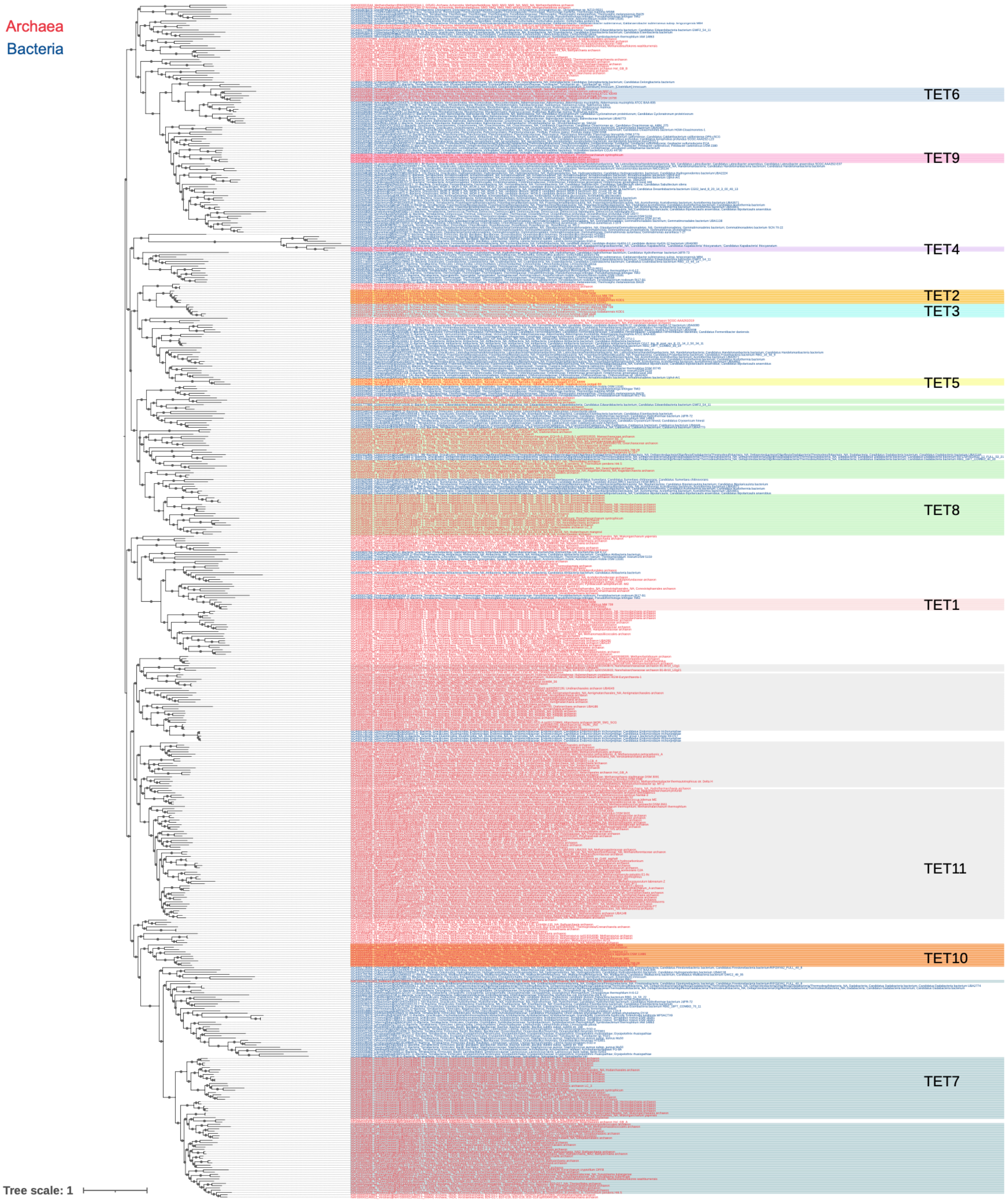

***Supplementary Fig. 6: (a) PsTETa AlphaFold 3 model*** *(ipTM score 0.87) featuring the novel insertion identified in some Asgard sequences, here colored in blue.* ***(b) Associated plDDT plot per residue.*** *The dashed red line indicates the confidence threshold (plDDT > 70), above which predicted structures are generally considered reliable. The region of the novel insertion (between residues 166 and 185) is highlighted in blue.*

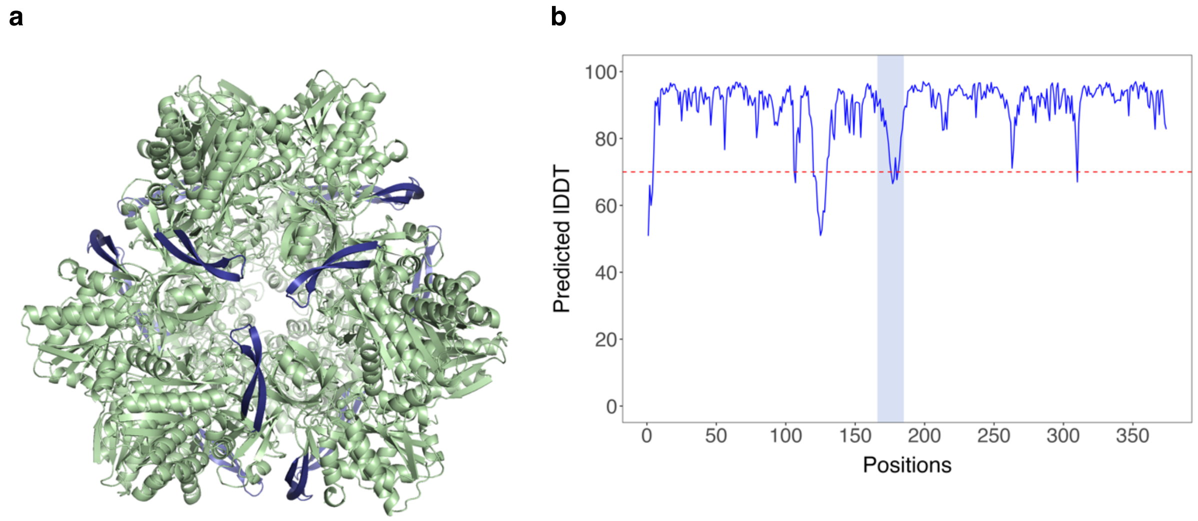
